# Inflammation is an essential trigger for POLE-driven ultramutated colorectal cancer

**DOI:** 10.64898/2026.10.19.760274

**Authors:** Md Kawsar Mustofa, Mitzi Aguilar, He Zhang, Debanjan Sarkar, Sambit Roy, Manzar Alam, Mohammad A. Basher, Subhransu S. Sahoo, Esra A. Akbay, Diego H. Castrillon, Hasan Zaki

## Abstract

Pathogenic variants in the exonuclease (proofreading) domain of DNA polymerase ε (*POLE*) are frequently detected in colorectal cancers (CRC) with high tumor mutational burden (TMB), yet their functional role in intestinal tumorigenesis remains unknown. Here, we investigated the impact of mutation in *POLE* on intestinal homeostasis using a conditional mouse model expressing the *Pole^P286R^* pathogenic driver in intestinal epithelial cells. *Pole^P286R^* mutant mice developed invasive adenocarcinomas in the small intestine with incomplete penetrance and extremely long latency but did not spontaneously develop colorectal tumors. However, induction of colitis with dextran sodium sulfate (DSS) resulted in a striking phenotypic shift to colorectal adenocarcinomas at high frequency. Molecular and histological analyses revealed hyperactivation of Wnt/β-catenin, NF-κB, ERK and STAT3 pathways, accompanied by markedly higher proliferation of intestinal epithelial cells, in DSS-treated *Pole^P286R^*mutant mouse colons. Whole genome sequencing of the tumor demonstrated exceptionally high TMB (>100 mutations/megabase) and characteristic trinucleotide signatures, consistent with human *POLE* tumors. Recurrent mutations in *Apc* and *Ctnnb1* underscored the central role of Wnt signaling in *POLE*-driven hypermutated CRC. In addition, intestinal organoids derived from *Pole^P286R^* mutant mice exhibited accelerated growth and increased expression of pro-proliferative genes. Together, these findings highlight POLE as a gatekeeper of intestinal genomic integrity and identify inflammation as the key driver that unleashes *POLE*-mutant CRC. Furthermore, this work establishes a mechanistically faithful preclinical platform for studying the molecular pathogenesis of CRC and advancing therapeutic drug discovery.

## INTRODUCTION

Colorectal cancer (CRC) is the third most common cancer and a leading cause of cancer-related deaths in the United States ^1^. More than 70% of CRC cases are sporadic, arising without any family history or inherited genetic predisposition and are driven by spontaneous accumulation of somatic mutations or epigenetic alterations ^2^. The risk of intestinal oncogenic modifications is influenced by several pathophysiological factors, including age, Western diet, metabolic disorders, dysbiosis, and chronic inflammation ^3^. Patients with inflammatory bowel disease (IBD), such as Crohn’s disease and ulcerative colitis, are at a significantly higher risk of developing CRC ^4^. Although the precise underlying mechanism of colitis-associated CRC remains poorly defined, inflammatory mediators are potential inducers of DNA damage and oncogenic mutations ^4–6^. In addition, the rapid regeneration of epithelial crypts in the injured colon imposes replication stress, further promoting replication errors ^7^. Somatic mutations in oncogenes and tumor suppressor genes ultimately drive neoplastic transformation of intestinal epithelium, leading to CRC development ^2, 8^.

Although most sporadic CRC display a relatively low tumor mutational burden (TMB), approximately 10-15% exhibit high TMB, defined by mutation rates ≥12 per megabase ^9^. Such “hypermutated” CRCs typically exhibit defective mismatch repair (MMR) or harbor pathogenic variants in *POLE* and *POLD1* ^2, 9^, which encode the housekeeping DNA replicases ε and δ, respectively ^10, 11^. Replication fidelity is impaired by specific somatic single amino acid substitutions in *POLE* or *POLD1*, with recurrent variants clustering in their proofreading domains, leading to accumulation of mutations. These pathogenic drivers produce “ultramutated” tumors with TMB typically >100/megabase. Among recurring *POLE*/*POLD1* pathogenic drivers, *POLE^P286R^* (p.Pro286Arg) is the most frequent and is associated with the highest TMB ^12^. Clinical relevance is underscored by polymerase proofreading-associated polyposis (PPAP), an autosomal dominant predisposition syndrome in which germline *POLE* or *POLD1* variants lead to early-onset colorectal adenomas and carcinomas with high TMBs ^13, 14^. *POLE*-driven ultramutation is most frequently observed in endometrial cancer (7–10%) and, to a lesser extent, in CRC (1-2%) ^15–17^.

Genetically engineered *Pole* mouse models have enabled in vivo study of polymerase-driven tumorigenesis. Whole body expression of *Pole^P286R^* resulted in high TMB tumors throughout the body ^18, 19^. In the conditional *LSL-Pole^P286R^* knock-in allele, a *loxP*-flanked transcriptional STOP cassette blocks *Pole^P286R^* expression until Cre-mediated excision^19^. Previously, we showed that combining *LSL-Pole^P286R^* with an endometrial Cre allele resulted in lethal endometrial cancers, with recapitulation of key aspects of human *POLE-driven* endometrial cancers ^20, 21^. In contrast, *POLE*/*POLD1* drivers are rare in lung cancer, and expression of *LSL-Pole^P286R^* alone in lung epithelium failed to induce tumors ^22^. However, when combined with *Kras^G12D^*, *Pole^P286R^* increased TMB, revealing striking tissue-specific susceptibilities to *POLE*-driven carcinogenesis ^22^. However, the functional contribution of *Pole^P286R^*in CRC has not yet been studied using a tractable, tissue-specific model.

In this study, we investigated the role of *POLE* in maintaining intestinal genomic integrity and host defense against colorectal neoplasia using a conditional mouse model heterozygous for *Pole^P286R^* in intestinal epithelial cells. Unlike endometrium-specific *Pole^P286R/+^*mice, intestinal-specific *Pole^P286R/+^*mice did not develop spontaneous colorectal tumors. Interestingly, however, when colitis was induced with dextran sodium sulfate (DSS), colorectal tumors developed with 100% penetrance. Whole genome sequencing (WGS) demonstrated exceptionally high TMB and characteristic trinucleotide signatures in those tumors. This study demonstrates that *POLE* exonuclease activity is indispensable for maintaining intestinal genomic stability under inflammatory stress. These findings provide significant insights into the mechanistic link between chronic intestinal inflammation and CRC and establish *LSL-Pole^P286R^* mice as a valuable preclinical model for dissecting molecular pathways underlying inflammation-driven colorectal carcinogenesis and testing CRC therapies.

## RESULTS

### Intestine-specific expression of *Pole^P286R^* leads to adenocarcinoma development in the small intestine but not colon

To generate intestinal epithelial cell-specific *Pole^P286R/+^* mice, we crossed *LSL-Pole^P286R/+^* mice with *Villin-Cre* mice, producing doubly-heterozygous *LSL-Pole^P286R/+^;Vil-Cre* experimental (hereafter referred to as PVC) and control (either *LSL-Pole^P286R/+^* or *Vil-Cre*) mice (Figure 1A). PVC mice co-express both wild-type and mutant *Pole* (*Pole^P286R^*) in intestinal epithelial cells, and only wild-type *Pole* in all other cells. Animals were born at expected Mendelian ratios and appeared healthy until adulthood. To assess if PVC mice spontaneously developed intestinal carcinogenesis, we initially monitored 7 PVC and 5 control mice. Three out of 7 PVC mice developed symptomatic disease, characterized by pallor and weight loss, necessitating euthanasia. Asymptomatic animals had normal hematocrits, but symptomatic animals were severely anemic with decreased hematocrits (Figure S1A). Reticulocyte counts were concordantly elevated, demonstrating that the anemia was likely the result of chronic blood loss and iron deficiency (Figure S1B).

**Figure 1.**
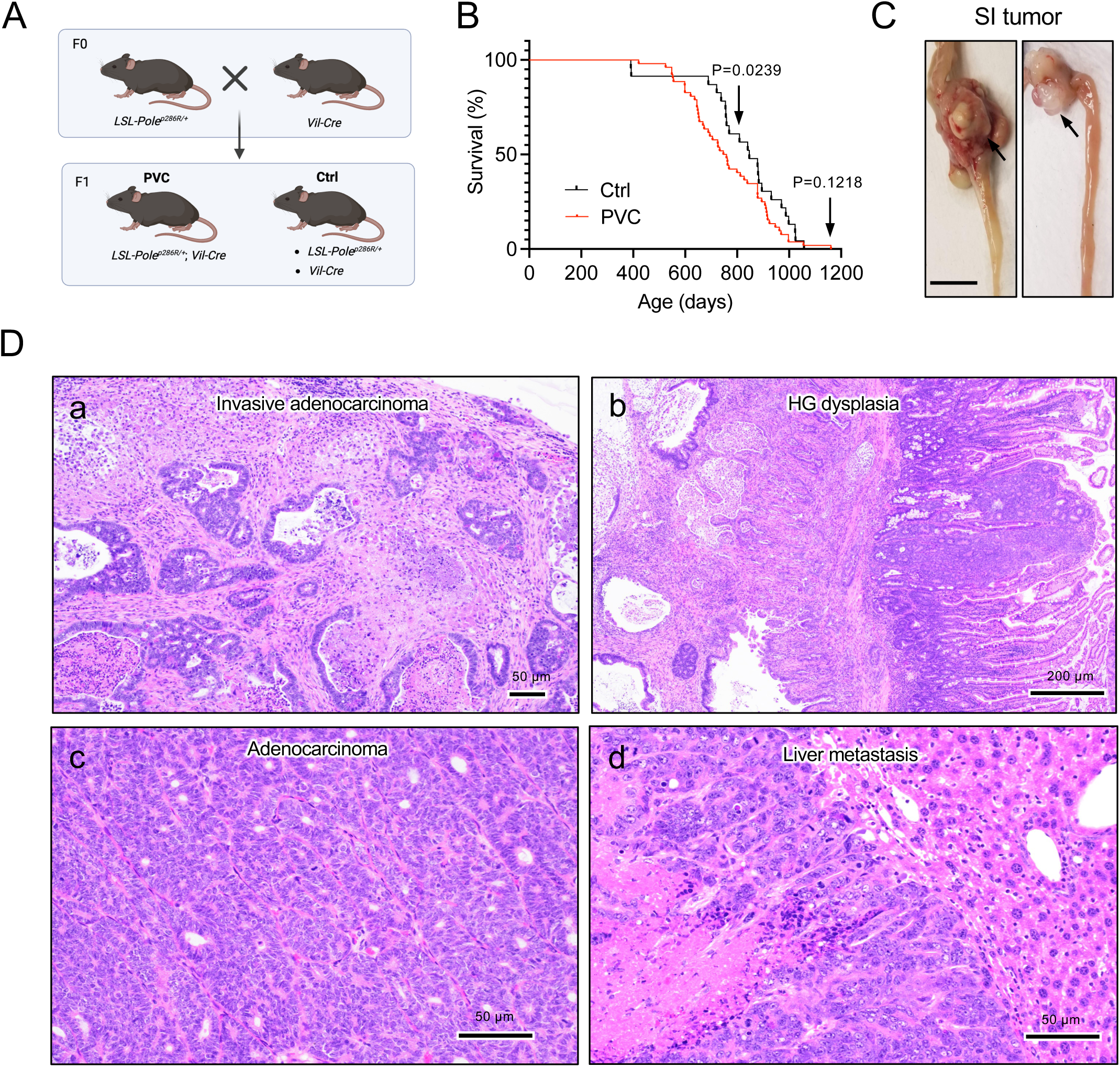
*Pole^P286R^* intestinal expression results in death and small intestinal neoplasia. (A) Breeding scheme for generation of intestinal epithelial cell-specific *Pole^P286R^* mutant mice, *LSL-Pole^P286R^/+;Vil-Cre* (PVC). Sibling control (Ctrl) and PVC mice are efficiently generated in a single generation. (B) Control and PVC mice was monitored for survival (*P*=0.1218 at end point but <0.0239 at 800 days). Statistical significance was determined by Log-rank test. (C) Symptomatic PVC animals were examined for intestinal tumor development. Symptomatic PVC mice harbored small intestinal neoplasms and metastases. Images show discrete small intestinal neoplasms in two different mice, bars=1 cm. (D) H&E-stained SI tumor sections from 4 different mice. (a) Well-differentiated invasive adenocarcinoma with transmural invasion to serosa. (b) Invasive adenocarcinoma in muscularis propria subjacent to mucosal high-grade (HG) dysplasia. (c) Moderately differentiated adenocarcinoma. (d) Liver with metastatic adenocarcinoma and necrosis.

Next, we followed the survival with a separate cohort of PVC (n=41) and control mice (n=21). The median survival was 753 days for PVC versus 846 days for controls (Figure 1B). While overall difference did not reach significance (p=0.1218, log-rank test), survival divergence became significant by 600 days (p<0.05) (Figure 1B). About one third of PVC mice exhibited tumor symptoms, necessitating euthanasia. There was no difference in body weight between control and PVC mice taken at endpoint (Figure S1C). At necropsy, all the symptomatic PVC mice (n=14) harbored isolated tumors in the small intestine, whereas no tumors were observed in the colon (Figure 1C). Histologic analysis of the small intestinal tumors revealed that they were well-differentiated invasive adenocarcinomas associated with intestinal intramucosal dysplasia (Figure 1D, a and b). In some animals, transmural invasion to serosa (Figure 1D, c) or metastases to the liver were present (Figure 1D, d). Collectively, unlike endometrium specific *Pole^P286R/+^* mice, intestinal epithelial cell-specific *Pole^P286R/+^* mice exhibited partial susceptibility to tumorigenesis and only in the small intestine. Adenocarcinomas in the small intestine showed aggressive features, albeit with incomplete penetrance and extremely long latency.

### Colitis triggers colorectal adenocarcinoma development in *Pole^P286R^*mutant mice

Absence of visible tumors in the colons of PVC mice was somewhat unexpected, given the high penetrance of endometrial cancer in the same genetic context and the occurrence of *POLE*/*POLD1* mutations in some colorectal cancers ^19, 21^. Unlike endometrial cancer, however, chronic inflammation is a well-established risk factor for CRC, and colitis-associated CRC frequently exhibits an ultramutator phenotype ^6, 23^. This prompted us to hypothesize that colitis might drive colorectal neoplasia in PVC mice. To test this, we induced colitis in PVC mice by administering 2.5% DSS for 5 days and repeating the treatment twice at 2-week intervals (Figure 2A). After completion of the third cycle of DSS, mice were maintained on regular drinking water and monitored for illness and survival. Surprisingly, the DSS-induced chronic colitis resulted in a significant acceleration of mortality in PVC mice, with >50% of animals requiring euthanasia by 300 days (Figure 2B) due to lethargy and rectal prolapse (Figure 2C). At necropsy, we observed extensive colorectal tumor development (Figure 2D), whereas untreated PVC mice developed no colorectal tumors.

**Figure 2.**
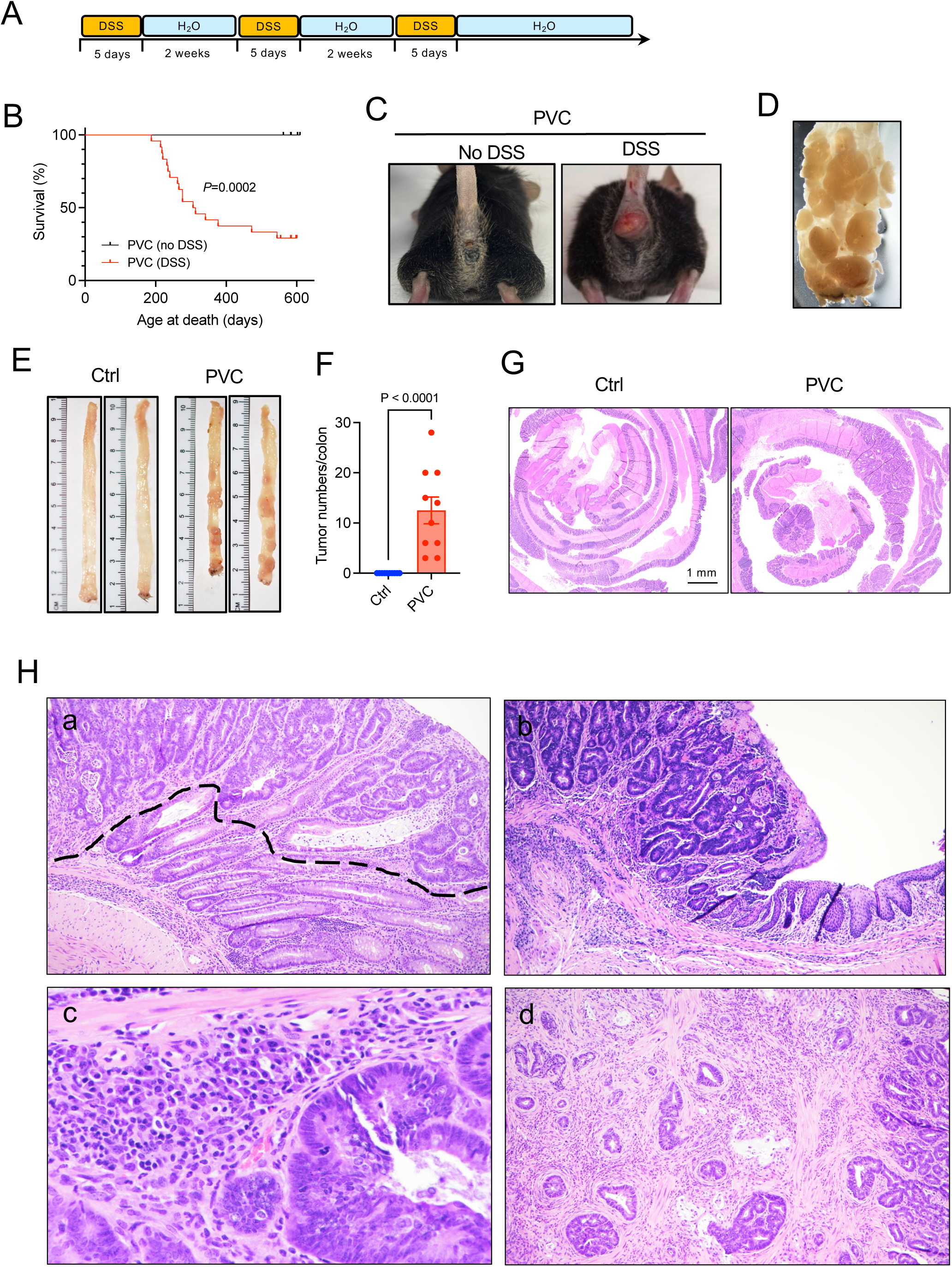
DSS administration drives colorectal carcinoma in PVC mice. (A) Model of DSS-induced chronic colitis. Mice were administered 2.5% DSS for 5 days, followed by regular drinking water for 2 weeks. The DSS cycles were repeated two more times. (B) Survival of DSS-fed and untreated PVC mice. *P* value per log-rank test. Animals with severe disease including rectal polyps were euthanized and scored as deaths. (C) Gross images of anorectal area of DSS-treated and untreated mice euthanized at 240 days. (D) A representative image of the rectum of a DSS-fed PVC mouse that developed rectal prolapse. (E-H) A separate group of control and PVC mice were treated with DSS regimen and euthanized at 12 weeks post-DSS. (E) Colons exhibiting multiple tumor nodules developed in PVC mice. (F) Tumor number per colon. Data represents mean ± SEM, *P* value per two-tailed unpaired t-test. (G) Colon rolls of H&E sections. (H) Histopathologic features of colorectal neoplasia in DSS-treated PVC mice. (a) Adenomatous (dysplastic) epithelium (above dashed line) overlying normal epithelium. (b) Adenomatous dysplasia extending to rectal squamocolumnar margin. (c) Example of abundant dense mixed infiltrate comprised of lymphocytes, plasma cells, and neutrophils. (d) Invasive adenocarcinoma infiltrating muscularis propria.

We then assessed pathological changes in the colons of PVC and control mice at a defined time point. All mice were euthanized 12 weeks after initial DSS administration. Interestingly, all PVC mice exhibited adenomatous polyps throughout the colon, while none of the DSS-fed control mice showed any visible tumors (Figure 2E and F). Tumors were more abundant in the mid and distal colon, ranging from 4-30 per mouse (Figure 2E and F). Histopathological examination confirmed neoplasia in PVC but not control colons (Figure 2G). Tumors of PVC mice were pathologically diverse with low-grade and high-grade dysplasia (Figure 2H, a). Adenomatous polyps occasionally developed in the anorectal area (Figure 2H, b), and were interspersed with immune cells comprised of lymphocytes, plasma cells, and neutrophils (Figure 2H, c). Notably, some PVC mice developed invasive adenocarcinoma infiltrating through the mucosa and into muscularis propria (Figure 2H, d), revealing an aggressive phenotype. These results demonstrate that *Pole^P286R^* mutation and inflammation synergize to a surprising degree to produce multiple independent colorectal adenocarcinomas in mice, shifting the neoplasia phenotype from the small to the large intestine.

### Colorectal tumors of PVC mice are characterized by increased expression of oncogenic and proliferative drivers and infiltration of immune cells

Colorectal tumorigenesis is frequently accompanied by aberrant expression of genes regulating proliferation, stemness and metastasis ^2^. To identify molecular signatures of tumors in DSS-fed PVC mice, we measured the expression of *Ctnnb1* (β-catenin)*, cMyc, Ccnd1* (Cyclin D1)*, Cdk4, Cdkn1a* (p21)*, Lgr5, Ki67* and *Mmp7* by real-time PCR. The expression of *Ctnnb1, cMyc, Ccnd1, Lgr5, Ki67, Cdk4*, and *Mmp7* was significantly elevated, whereas *p21* was reduced compared to control and non-tumor regions of PVC colons (Figure 3A). Western blot analysis and immunohistochemistry confirmed that cMyc, Cyclin D1, and β-catenin expression were upregulated in PVC tumors (Figure 3B-D). In agreement with higher expression of the pro-mitotic factors, the proliferation marker PCNA was strongly expressed in PVC mouse tumors (Figure 3B-D).

**Figure 3.**
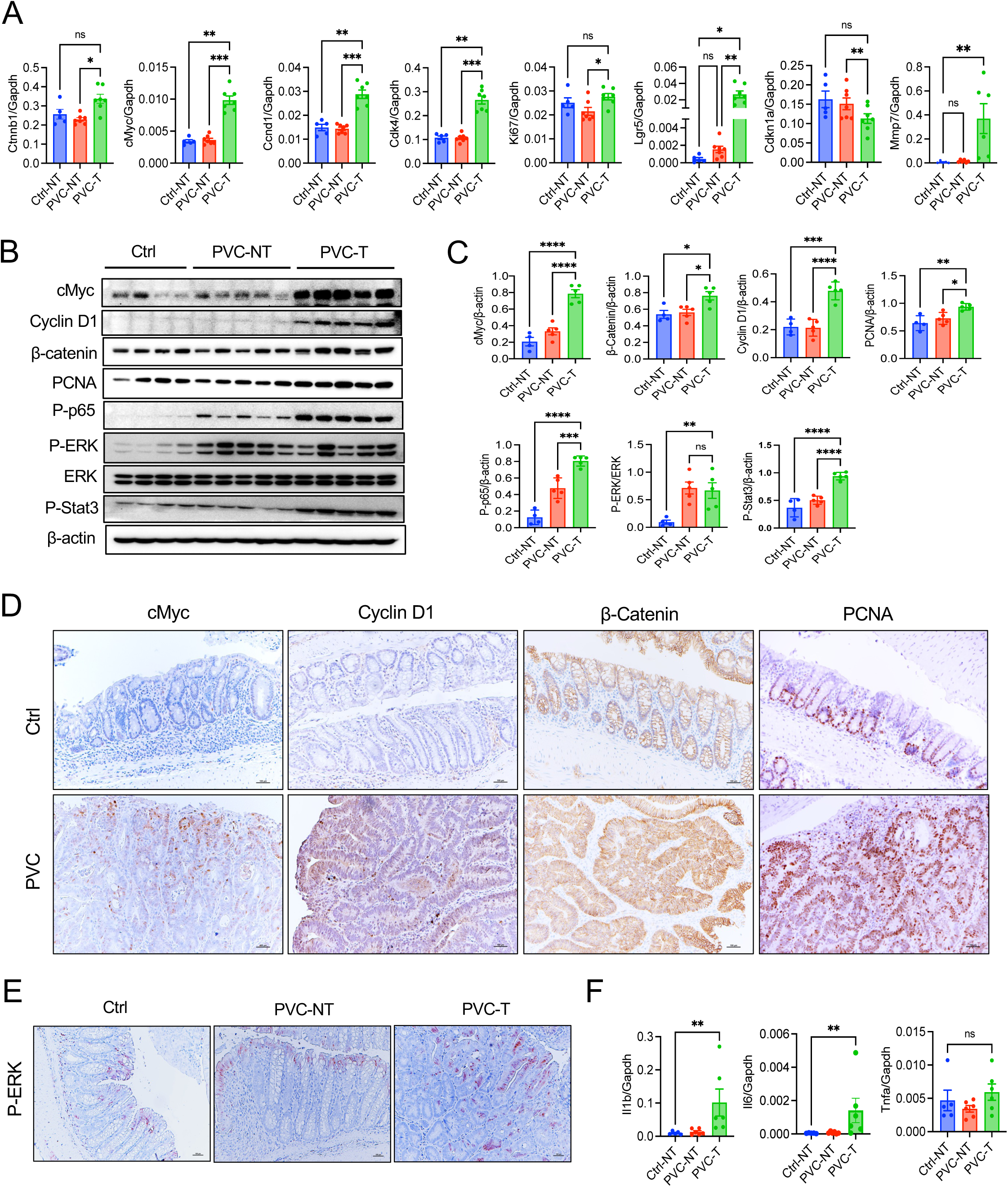
Tumor induction of PVC mice is associated with increased expression of proliferative genes. Chronic colitis was induced in control and PVC mice following protocol shown in Figure 2A. Colorectal tumor development was assessed at 12 weeks post colitis induction. (A) RNA isolated from the colon of control and PVC mice were measured for indicated genes by real-time RT-PCR. NT: non-tumor; T: tumor. Data represents means ± SEM, *P<0.05, **<0.01, ***P<0.001, ****P<0.001 using one-way ANOVA test. (B) Tissue homogenates were analyzed for the expression of cMyc, Cyclin D1, β-catenin, PCNA and activation of p65, ERK, and STAT3. (C) Densitometric analysis of immunoreactive bands shown in B. (D) Colon sections were immunostained for cMyc, Cyclin D1, β-catenin and PCNA. (E) Immunostaining of P-ERK in control and PVC mouse colons. (F) Expression of *Il1b*, *Il6* and *Tnfa* measured by real-time PCR. Data represents means ± SEM, *P<0.05, **<0.01, ***P<0.001, ****P<0.001 per one-way ANOVA test.

Several inflammatory signaling pathways, including NF-κB, ERK and STAT3, promote epithelial cell proliferation and are upregulated in CRC ^5^. We analyzed the activation of these pathways by western blotting, revealing NF-κB, ERK, and STAT3 hyperactivation in PVC mouse tumors as compared to DSS-treated control mouse colons (Figure 3B and C). Immunohistochemistry confirmed ERK pathway activation in tumor epithelium of PVC mice (Figure 3E). Consistent with higher activation of inflammatory pathways, proinflammatory cytokines IL1β and IL6 were markedly upregulated (Figure 3F).

To gain insight into the immune microenvironment of PVC tumors, we evaluated colons 12-weeks post-DSS treatment with F4/80 and CD3, which label macrophages and T cells, respectively. Both control and PVC mouse colons showed macrophage and T cell infiltration into the lamina propria and submucosa (Figure S2). Tumors of PVC mice were also infiltrated with macrophages to a similar extent as non-tumorous colonic mucosa (Figure S2). Interestingly, adenomas of PVC mice were infiltrated with extensive CD3^+^ T cells (Figure S2). Non-tumor regions of PVC and control colons exhibited comparable levels of CD3^+^ T cell infiltration (Figure S2). Notably, T cell abundance was lower in the stroma of adenocarcinoma than hyperplastic and low-grade dysplastic lesions (Figure S2). Collectively, these data suggest that *Pole^P286R^* allele drives neoplastic alterations in the colonic epithelium under the influence of inflammation, leading to upregulation of proliferative and oncogenic signaling. Furthermore, *Pole^P286R^* mutant colorectal tumors exhibited immunogenic features characterized by extensive T cell infiltration.

### Tumors of PVC mice harbor high mutational burdens and characteristic *POLE* trinucleotide signatures

CRC development in PVC mice following colitis induction suggests that inflammation synergizes with the oncogenic mutations resulting from aberrant Pole exonuclease activity. To identify recurrently mutated genes and TMB in the context of *P286R* mutation, we performed WGS of both small intestinal tumors from untreated PVC mice (n=2) and large intestinal tumors from DSS-treated PVC mice (n=8).

Both types of specimens yielded extremely high TMBs, much higher than described in prior intestinal tumor models and in line with human ultramutated cancers and *Pole* endometrial and other cancer models ^19, 21, 22, 24^ (Figure 4A). The higher TMB of 800 per megabase (Mb) in colorectal vs. 90 per Mb in small intestinal tumors is attributable due to the presence of multiple confluent tumors in the segments of colon used for DNA preparation (Figure 4A), whereas the small intestinal tumors were clearly singular. Tumors harbored expected trinucleotide signatures with a predilection of T-G substitutions. There was a highly reproducible shift towards C>A, C>T, T>C, and T>G substitutions; however, the signatures were slightly altered in the small intestinal vs. colorectal tumors (Figure 4B). These point mutations are predicted to result in large numbers of single amino acid substitutions (missense mutations), and also abundant premature stop (frameshift), splicing, and read-through mutations, the latter especially likely to generate neoepitopes (Figure 4C). As expected, given the base substitution phenotype associated with *Pole^P286R^*, indels were relatively rare. Tumors also did not exhibit large-scale chromosomal deletions or amplifications (Figure 4D).

**Figure 4.**
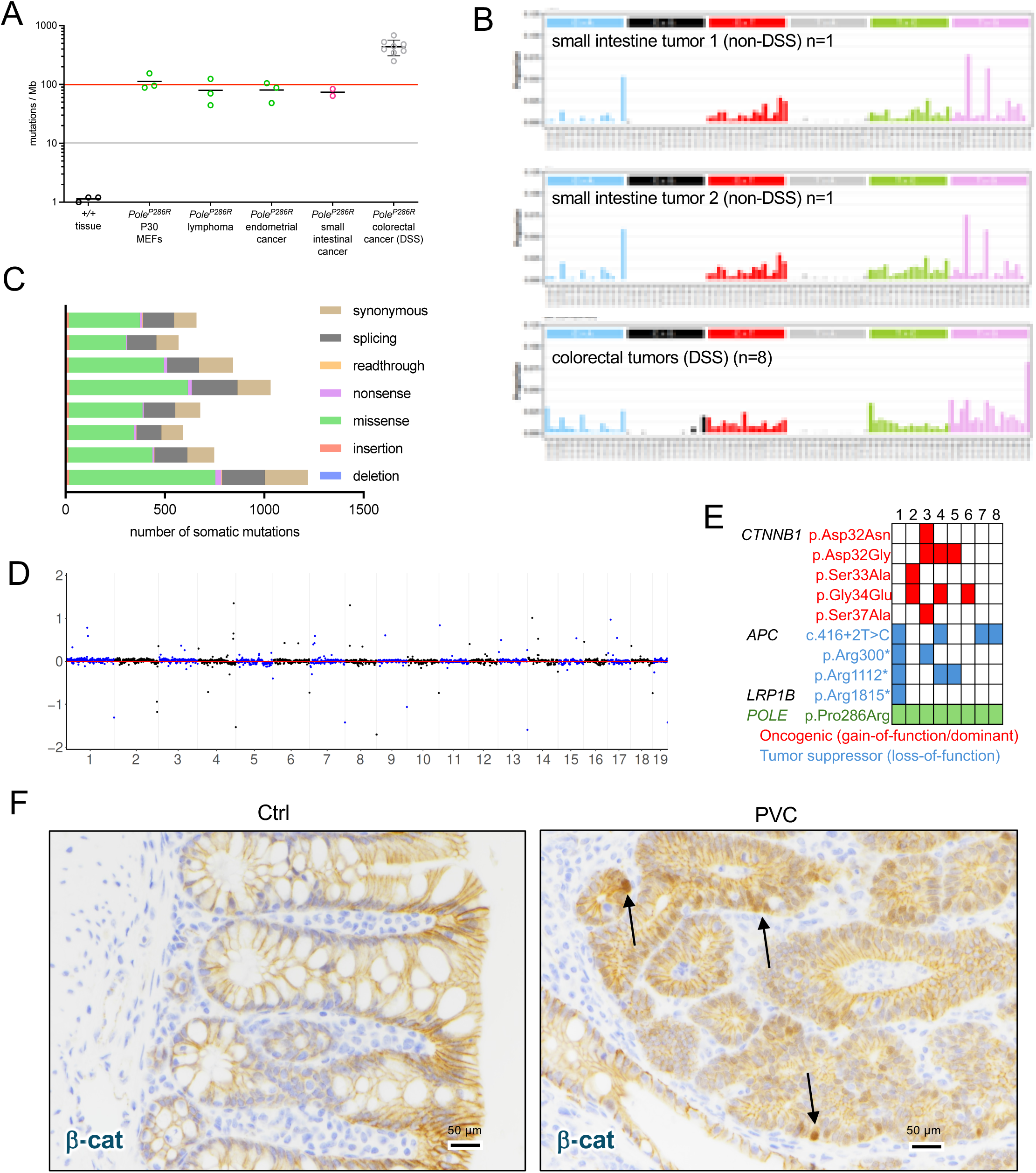
*Pole^P286R^*tumors in PVC mice display high TMB. Small intestinal tumors from 2 non-DSS treated mice and colorectal tumors from 8 mice (12 weeks post DSS) were subjected to WGS. (A) TMB, mutations per Mb. Data for passage 30 mouse embryo fibroblasts (P30 MEFs), lymphomas, and endometrial cancers is from prior publications ^21, 24^. (B) Trinucleotide mutational signatures from two independent small intestinal tumors and n=8 colorectal tumors in aggregate. (C) Impact of single nucleotide substitutions (mutation types). Somatic mutations located within coding sequences were assigned to different categories using the SnpEff tool with GENECODE M16 as a reference. The x-axis represents the number of somatic mutations in each sample. (D) Copy number plots from WGS data. The y-axis shows the normalized depth (log_2_(depth/median depth across genome)); x-axis represents position among the 19 autosomes as illustrated in legend. (E) Mutations in canonical intestinal cancer genes including recurring mutations in *Ctnnb1* (β-catenin) and *Apc*. Detection of >1 mutant allele is consistent with presence of multiple independent tumors in colorectal segments used for DNA isolation. (F) β-catenin immunohistochemistry of control and PVC colons, showing its localization. Focal nuclear localization was detected in PVC tumors.

Analyses of canonical intestinal cancer drivers demonstrated recurring mutations in *Ctnnb1*, *Apc*, and *Lrp1b*, with *Pole^P286R^* detectable in all samples, as expected. (Figure 4E). The *Ctnnb1* mutations were all canonical exon 3 point mutations that are known recurring hotspots leading to β-catenin stabilization/nuclear localization, and all specimens harbored either a *Ctnnb1* or *Apc* mutation (Figure 4E). Immunohistochemistry confirmed nuclear localization of β-catenin in PVC tumors (Figure 4F). These results demonstrate that the Wnt/β-catenin pathway is a major target in *Pole^P286R^*-driven CRCs, consistent with prior results showing that *Pole*-mediated tumorigenesis is dependent upon selection for misregulation of canonical cancer pathways.

### Increased epithelial proliferation following acute colitis leads to faster resolution of mucosal damage and development of dysplastic lesions in PVC mice

We next investigated whether *P286R* mutation impacts pathogenesis and resolution of colitis. Both control and PVC mice were administered 2.5% DSS for 5 days to induce acute colitis (Figure 5A). We monitored body weight, diarrhea scores and occult bleeding during acute colitis. PVC and control mice exhibited comparable clinical signs of colitis (Figure 5B-D). After 14 days of recovery period post completion of DSS treatment, mice were sacrificed, and colon lengths were measured. However, there was no significant difference between the groups, indicating similar colitis susceptibility for control and PVC mice (Figure 5E and F).

**Figure 5.**
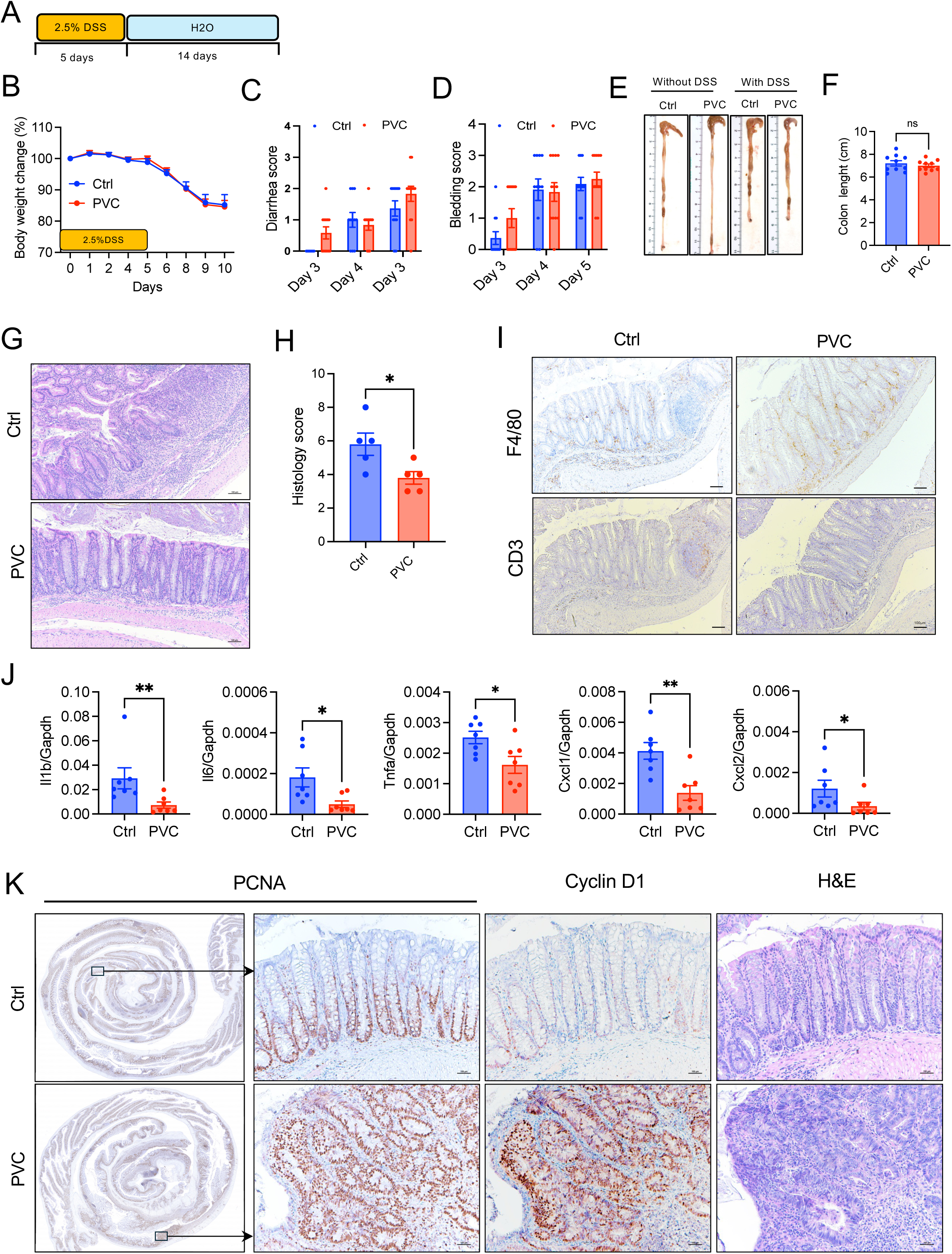
PVC mice exhibit faster recovery from DSS-induced colitis. (A) Control and PVC mice were administered 2.5% DSS for 5 days. Following 14 days of recovery mice were sacrificed. Body weights (B), diarrhea (C) and rectal bleeding (D) were scored. (E) Gross image of colons from DSS-untreated and treated mice. (F) Colon lengths of DSS-fed control and PVC mice. (G) Representative H&E-stained colon sections. (H) Combined histological scores for inflammation, ulceration and crypt destruction. (I) Colon tissue sections showing immunostaining of macrophages (F4/80) and T cells (CD3). (J) Real-time RT-PCR analysis of proinflammatory cytokines and chemokines. (K) Colon tissue sections immunostained for PCNA and Cyclin D1, and stained with H&E. Data represents means ± SEM, *P<0.05, **<0.01, two-tailed unpaired t-test.

Histopathological assessment of inflammation, mucosal ulceration, and crypt destruction were then evaluated (Figure 5G and H). Interestingly, control colons exhibited greater infiltration of inflammatory cells in the submucosa and lamina propria, with loss of crypt structure as compared to PVC colons (Figure 5G). Consistently, there was increased macrophage and CD3^+^ T cell infiltrates into the lamina propria and submucosa of control colons (Figure 5I). Furthermore, the expression of proinflammatory cytokines and chemokines, including *Il1b*, *Il6, Cxcl1,* and *Cxcl2*, were significantly reduced in PVC mouse colons compared to those in control mice (Figure 5J). Despite equal body weight loss and clinical signs of colitis, reduced inflammation at day 14 post DSS cycle in PVC mice suggests that recovery and healing of DSS-induced colonic damage were faster in PVC mice.

During acute injury or colitis, intestinal epithelial cells rapidly proliferate to replace the injured mucosa and maintain epithelial barrier integrity. To evaluate the impact of pathogenic *Pole^P286R^*driver on the proliferative response of colonic epithelium during the recovery stage of DSS-induced colitis, colon sections collected at day 14 post DSS treatment were stained for PCNA. The overall number of PCNA-positive cells per crypt was higher in PVC mice (Figure S3A and B). We further examined the active proliferation by BrdU incorporation assay and observed increased BrdU-positive cells in PVC colons (Figure S3C and D). Notably, localized regions displayed strong immunoreactivity for PCNA (Figure 5K). Histological examination revealed that those densely PCNA-positive regions represent minute foci of dysplasia (Figure 5K and S2E). The dysplastic areas also strongly expressed Cyclin D1 (Figure 5K), suggestive of early neoplastic transformation. Together, these findings demonstrate that *Pole^P286R^* mutation does not enhance acute colitis susceptibility but rather enhances host response for healing and inflammation resolution. Furthermore, early dysplastic changes occur shortly after colonic injury in PVC mice, most likely due to defective DNA replication fidelity, increased TMB, and selection for cancer-driving mutations.

### Tumorigenesis in *Pole^L/+^;Vil-Cre* mice is proportional to intestinal inflammation

Given our observation of dysplastic lesions as early as 14 days post-DSS (Figure 5K), we wondered whether a single cycle of DSS would be sufficient to initiate tumorigenesis in PVC mice. To address this, we induced colitis in *Pole^L/+^;Vil-Cre* and control mice with 2.5% DSS for 5 days, followed by resting on normal drinking water. Mice were sacrificed at 4 and 12 weeks after withdrawal of DSS to document colonic tumor burden (Figure 6A). Remarkably, 60% of PVC mice developed visible colonic polyps within 4 weeks, and 100% developed tumors by 12 weeks (Figure 6B and C). Histopathological examination confirmed the development of hyperplasia, low-grade dysplasia, and high-grade dysplasia at 4 weeks post DSS in the colons of PVC mice (Figure 6D). Consistent with these phenotypic changes and prior results, there was increased expression of the proliferative markers *cMyc, Ccnd1, Ki67* and *Lgr5* in the same tissue samples (Figure 6E). Western blotting and immunohistochemistry further demonstrated increased levels of cMyc, Cyclin D1, and P-ERK in PVC tumors at 4 weeks post DSS treatment (Figure 6F and Figure S4).

**Figure 6.**
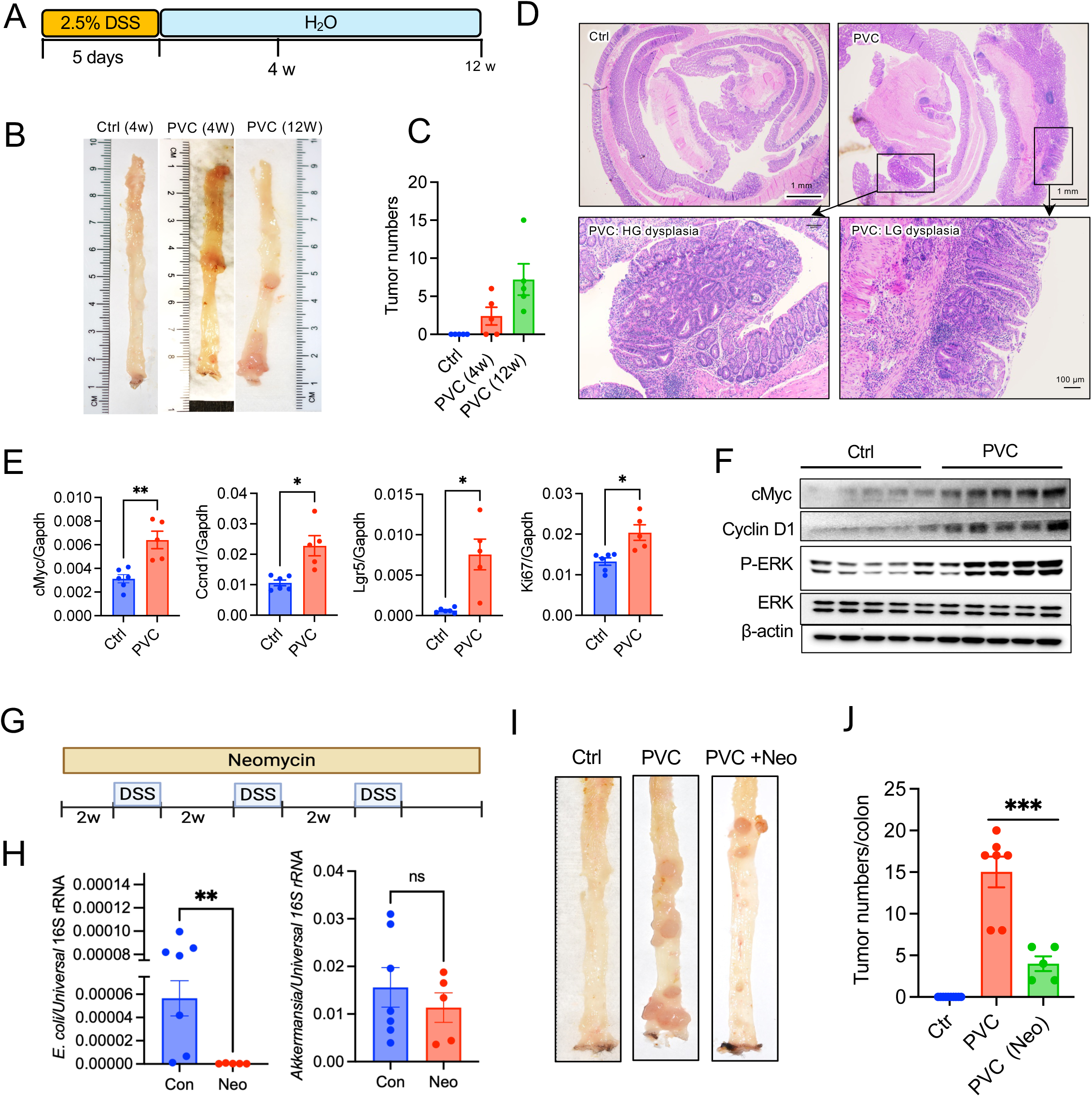
Colorectal tumorigenesis of PVC mice is correlated with the severity of DSS-induced colitis. (A) Control and PVC mice were administered 2.5% DSS for 5 days. Tumor development was monitored at 4- and 12-weeks post DSS. (B) Gross images of colons 4- and 12-weeks post DSS. (C) Visible adenomas polyps were counted. (D) H&E-stained colons collected at 4 weeks post DSS, showing development of hyperplasia, low-grade dysplasia and high-grade dysplasia in PVC mice. Scale bar=100μm. (E) Expression of indicated genes in control and PVC mouse colons at 4 weeks post DSS. (F) Colon tissues lysates were measured for the expression of cMyc, and Cyclin D1, and activation of ERK by Western blotting. (G) Chronic colitis was induced in control and *Pole^L/+^;Vil-Cre^/+^*mice with 3 cycle of 2.5% DSS (5 days), with 2 weeks interval between the cycles. A group of *Pole^L/+^;Vil-Cre^/+^* mice were treated with neomycin throughout the study. (H) Fecal DNA isolated from 4 weeks post neomycin treatment were measured for E. coli and Akkermansia with real-time qPCR. (I) Colons were collected 12 weeks post DSS treatment were examined for tumor development. (J) The tumor burden was counted. Data represents means ± SEM, ***P<0.001, two-tailed unpaired t-test.

Colorectal adenoma development in four weeks following one cycle of 2.5% DSS administration is remarkable as no genetic mouse models to date have developed CRC in such a short period. To verify if tumorigenesis can be further accelerated, we induced more severe colitis with the administration of 3% DSS for 7 days (Figure S5A). This increased colitis insult accelerated tumor induction and progression as evidenced by visible adenoma development with 100% penetrance within 4 weeks (Figure S5B and C). Histopathological examination revealed that tumors progressed to high-grade dysplasia and adenocarcinoma (Figure S5D). These findings imply that even a single episode of acute colitis is sufficient to induce colorectal adenocarcinoma in PVC mice.

To further test whether tumorigenesis is driven by inflammation per se rather than DSS exposure, we treated DSS-fed mice with antibiotic neomycin, (Figure 6A). Neomycin treatment markedly reduced *E. coli* burden, but not all gram-negative bacteria; for example, Akkermansia was not affected by neomycin (Figure 6H). Interestingly, neomycin-treated *PVC* mice exhibited significantly fewer tumor burden compared to antibiotic untreated *PVC* mice (Figure 6I and J). Together, these data suggest that inflammation is an essential driver for colorectal tumorigenesis in PVC mice and the oncogenic process is proportional to the severity of colitis.

### PVC mouse-derived intestinal organoids are hyperproliferative

To determine whether the TMB and specific driver mutations identified by WGS following DSS-induced colitis confer enhanced cellular fitness and proliferative capacity, we cultured colonic crypts from DSS-treated and untreated PVC mice using a 3D organoid system. Colorectal tumors were induced in PVC mice with the DSS regimen as shown in Figure 2A. Crypts were isolated from tumor-bearing colonic mucosa or DSS-untreated healthy colons of PVC mice and cultured for organoid development. Spheroids from tumor-bearing PVC colons grew significantly larger than those from normal mucosa of DSS-untreated PVC mice (Figure 7A and B) and expressed significantly higher *cMyc*, *Ccnd1*, *Ctnnb1*, and *Lgr5* (Figure 7C). Consistent with enhanced cell proliferation, immunofluorescence staining showed strong immunoreactivity for β-catenin in tumor-derived organoids, indicating aberrant activation of the Wnt/β-catenin pathway (Figure 7D).

**Figure 7.**
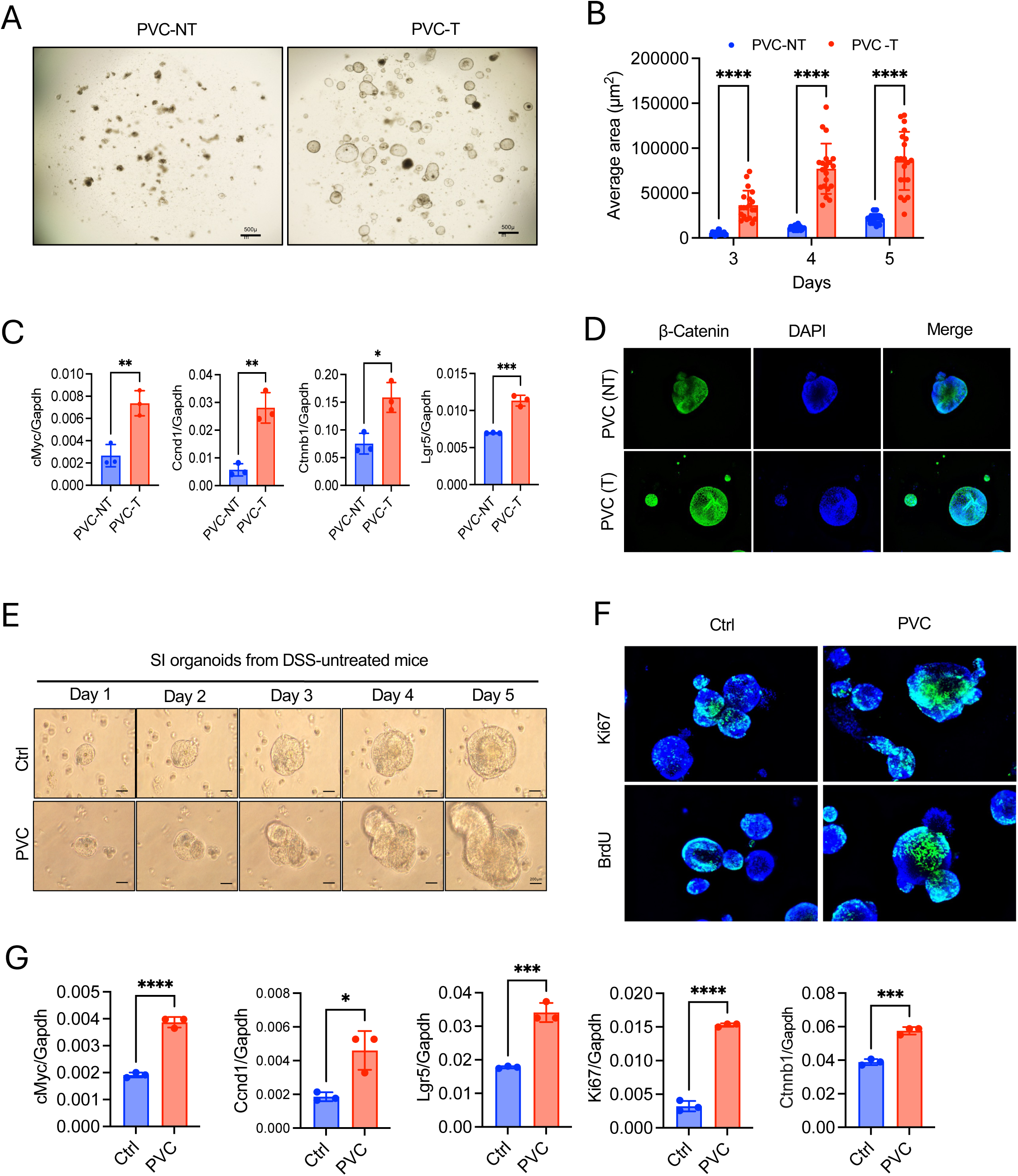
Intestinal organoids from PVC mice tumor are hyperproliferative. PVC mice were treated with 3 cycles of 2.5% DSS as shown in Figure 2A. Crypts isolated form tumors and DSS-untreated colons of PVC mice were cultured in a 3D organoid culture system. (A) Images of spheroids at day 4. (B) The size of the spheroids was measured at day 3, 4 and 5. (C) The expression of *cMyc, Ccnd1, Ctnnb1,* and *Lgr5* was measured by real-time RT-PCR. (D) Organoids were immunostained for β-catenin (green), nuclei counterstained with DAPI (blue). (E-H) Small intestinal crypts from healthy (DSS-untreated) control and PVC mice were cultured. (E) Organoid growth was monitored from day 1 to day 5. (F) Proliferation was assessed by Ki67 or BrdU staining (Green). Nuclei were counterstained with DAPI (blue). (G) RNA isolated at day 5 were measured for *cMyc, Ccnd1, Ctnnb1, Ki67* and *Lgr5*. Data represents means ± SD, *P<0.05, **<0.01, ***P<0.001, ****P<0.001, two-tailed unpaired t-test.

Next, we interrogated whether *Pole^P286R^* mutation affects innate proliferation and growth properties of intestinal epithelial cells. To address this, we cultured small intestinal (SI) and colonic crypts isolated from DSS-untreated control and PVC mouse colons. Surprisingly, both SI and colonic organoids from DSS-untreated PVC mice displayed faster growth than those of control mice (Figure 7E; Figure S6A and B). Ki67 and BrdU immunostaining further confirmed SI organoids of PVC mice have higher proliferative indices (Figure 7F). Concordantly, there was higher expression of *cMyc*, *Ccnd1*, *Ki67*, *Ctnnb1* and *Lgr5* in SI organoids of PVC mice (Figure 7G), suggesting that even without inflammatory insult, mutations may accumulate during replication cycles in intestinal epithelium of PVC mice, providing their proliferative fitness.

Collectively, our in vivo and in vitro findings demonstrate that *Pole^P286R/+^*impairs replication fidelity in intestinal epithelial cells, particularly under condition of replication, inflammatory and oxidative stress. This defect promotes hyperproliferation through the accumulation of oncogenic mutations and activation of proliferative signaling pathways (Figure 8).

**Figure 8.**
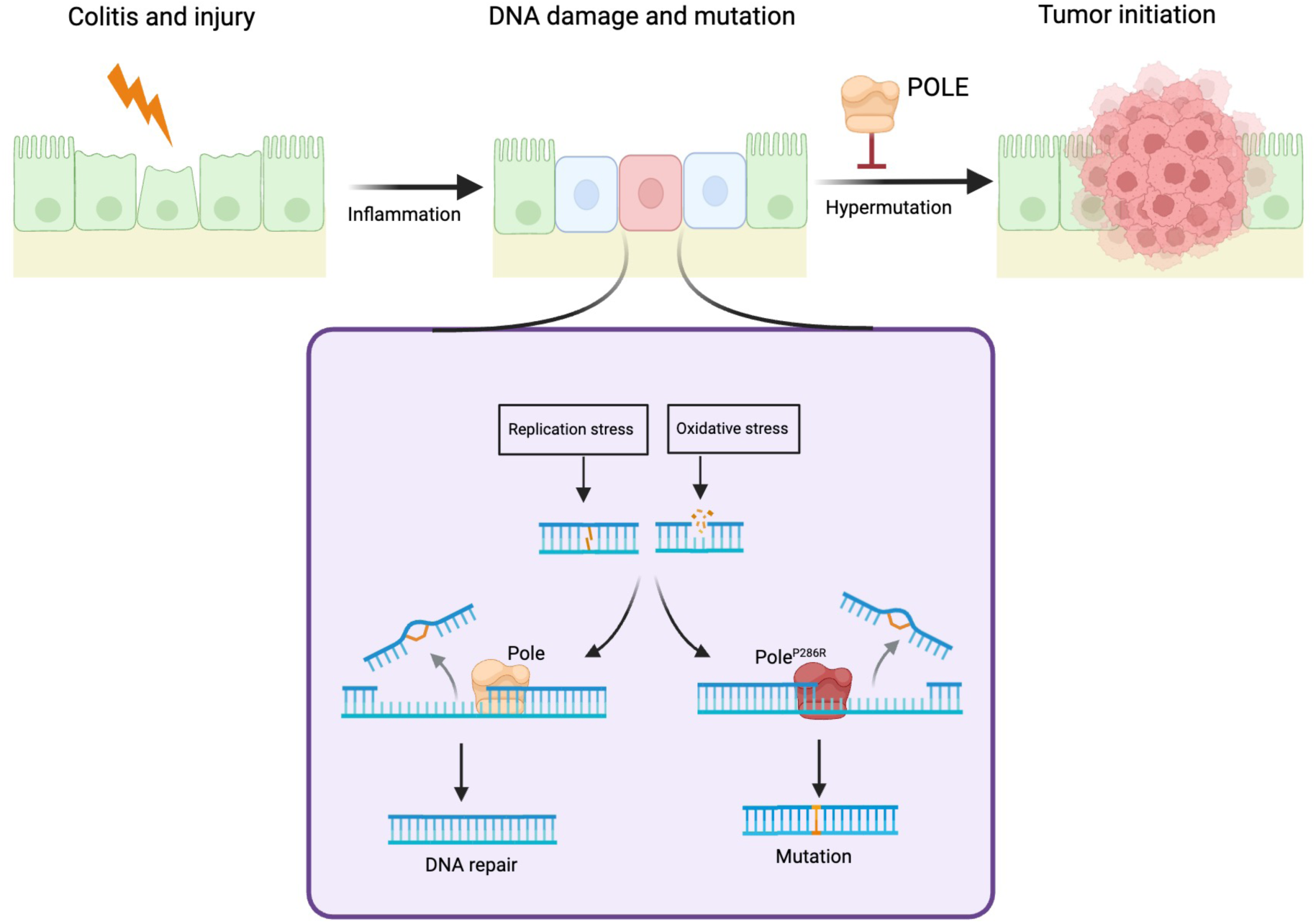
Proposed mechanism of inflammation-induced CRC development in Pole mutant mice. Intestinal injury and inflammation during colitis leads to DNA damage and mutations due to proliferative and oxidative stress. POLE plays an essential role in repairing DNA damage and mutations, maintaining genomic integrity. When proof reading function of POLE is dysfunctional, mutations accumulate in epithelial cells of inflamed colon, resulting in neoplastic transition and initiation of colorectal adenoma.

## DISCUSSION

CRC is a heterogenous disease that develops through diverse genetic and epigenetic alterations ^8^. While approximately 80% of CRC cases are non-hypermutated and driven by inherited or sporadic mutations in canonical oncogenes or tumor suppressor genes, more than 15% of CRCs display a high TMB, encompassing mutations in both driver and passenger genes ^25^. Notably, *POLE* oncogenic drivers have been identified in a subset of CRC with high TMB ^9, 17, 26, 27^. In this study, we provide functional evidence that the pathogenic *Pole^P286R^* mutation, which is the initiating oncogenic event in a subset of CRC and other cancers, predisposes mice to intestinal tumorigenesis. We demonstrate that intestinal injury and inflammation synergize with *Pole^P286R^* to dramatically increase mutational burden and promote neoplastic transformation.

Our previous work showed that endometrium-specific *Pole^P286R/+^*mice develop endometrial carcinoma with 100% penetrance ^21^. However, intestinal epithelial cell-specific *Pole^P286R/+^* mice showed only partial susceptibility, developing adenocarcinomas restricted to the small intestine. Similarly, mice expressing *Pole^P286R^* in the lung did not spontaneously develop lung adenocarcinoma ^22^. We and others have also observed that *Pole* and *Pold1* mutant (whole body) mice commonly developed tumors in endometrium, skin, and lymphoid tissues, but not in the colon ^24, 28^. These differences in cancer susceptibility underscore the tissue-specific role of *POLE.* Despite intensive investigation, the remarkably variable cancer susceptibility of diverse tissues to *POLE/POLD1*-driven tumorigenesis remain unexplained. However, distinctive and highly recurrent base substitution patterns evidenced by highly recurrent trinucleotide signatures may favor specific combinations of oncogenic driver events likely to result in tumors in specific contexts.”

The most striking observation in this study is that *Pole^P286R^* mutation renders mice highly susceptible to CRC only when they were exposed to a colitis-inducing inflammatory trigger. 100% of PVC mice developed colorectal tumors following a single cycle of DSS treatment, with tumors detectable as early as one month post colitis induction. This observation is clinically very significant since IBD is a major risk factor for CRC ^5, 6^. Conventional mouse models rely on chemical carcinogens, such as azyoxymethane, or mutations in oncogenes like *Apc* or *Kras* to induce colorectal tumors ^29, 30^. While these models are useful, the oncogenic mutations confound efforts to dissect the contribution of inflammation alone in mutagenesis and tumorigenesis. Since *Pole^P286R^* favors accumulation of oncogenic driver mutations under inflammatory stress, PVC mice are a powerful model to study how inflammation drives tumorigenesis. Our data demonstrate that a single cycle of DSS is sufficient to induce substantial tumor burden in PVC mice, establishing a direct and potent causal effect of intestinal inflammation in inducing tumorigenesis. This study helps explain why a large percentage of IBD-associated CRCs exhibit mutations in MMR factors or *POLE/POLD1* ^6, 23^. WGS confirmed that *Pole*-mutant colorectal tumors exhibit very high TMB ^21^. We identified recurrent clinically relevant mutations in *Apc* and *Ctnnb1*. Mutations in *Tp53*, *Kras*, and *Braf* were absent, but the number of tumors analyzed was small, preventing definitive interpretations. The most frequently mutated genes identified in our WGS were *Apc* and *Ctnnb1*, two well-established CRC drivers that lead to deregulated Wnt signaling. ^9, 31^. Interestingly, we also identified a truncating mutation in *Lrp1b*, a common early driver of CRC ^31^. *Lrp1b* was also identified as the most commonly mutated gene in intestinal tumors in whole-body *Pole* mutant mice ^28^. However, the precise role of Lrp1mutation in CRC remains to be explored.

It is well known that enteric pathogens, such as *Fusobacterium nucleatum*, *Escherichia coli*, and *Bacteroides fragilis*, are associated with increased CRC susceptibility ^32, 33^ ^34^. These bacteria can secrete genotoxins that directly induce DNA damage, thereby increasing mutational load, or promote tumorigenesis indirectly by driving chronic inflammation ^35–37^. Our data showing that depletion of gram-negative bacteria, particularly Enterobacteriaceae, with neomycin markedly reduced tumor burden further supports the role of gut microbiota in colitis and CRC pathogenesis ^32, 38^. Notably, we used neomycin alone instead of a cocktail of broad-spectrum antibiotics (ampicillin, neomycin, vancomycin and metronidazole) because complete depletion of gut microbiota renders mice highly susceptible to DSS-induced colitis due to impaired epithelial regeneration ^39, 40^. This likely explains why only a partial reduction in tumorigenesis was observed as neomycin does not eliminate all colitis causing bacteria. Nevertheless, our findings suggest that under pathological conditions, gut pathobionts contribute to mutational accumulation within colonic epithelial cells, while POLE proofreading activity serves as an essential protective mechanism to mitigate such mutation-driven oncogenic initiation.

Animal models of intestinal cancer are well-established tools to recapitulate human tumor biology and test new hypotheses or therapies. While existing genetic and chemically-induced models have been remarkably useful for studies of oncogenic signaling pathways, TMB is far below human counterparts, pointing to the need for new approaches to optimize intestinal cancer models with respect to TMB, which produces mutation-associated neoantigens and defines many aspects of tumor biology, clinical behavior, and treatment response ^41–43^. Mouse models based on MMR deficiency have been used to study the intersection of TMB, immunosurveillance and ICB, but do not yield sufficiently high TMB to fully elicit tumor immunogenicity ^24, 44, 45^. Compared with *Apc^Min^* models, in which tumors arise in the small intestine, our model faithfully mirrors human colitis-associated CRC. Importantly, *Pole^P286R^*-driven tumors exhibit exceptionally high TMB, producing mutation-associated neoantigens relevant for clinical behavior and treatment response ^46^. Thus, PVC mice provide a powerful system to study molecular mechanisms of hypermutated CRC and colitis-associated CRC, and to test the efficacy of cancer therapies.

In summary, this study identifies pathogenic *POLE* mutations as a critical driver of inflammation-induced colorectal tumorigenesis. Mice expressing intestinal epithelial cell-specific *Pole^R286R^* represent a unique and tractable preclinical model for investigating pathogenesis of hypermutated CRC and human colitis-associated CRC and should provide a robust platform for drug discovery. Our model requires only a single genetic cross, followed by a single DSS treatment to induce colitis and yield CRC with high penetrance, greatly facilitating and simplifying experimental protocols. Moreover, our findings demonstrate that *POLE* mutation is a potent oncogenic driver in an inflamed gut and suggest that genetic screening for *POLE* mutations in high-risk IBD patients, combined with strategies to modulate inflammation, may provide effective strategies for CRC prevention and treatment.

## METHODS AND MATERIALS

### Genetically engineered mouse model

Mice were housed in a pathogen-free animal facility and fed ad libitum on standard chow. Both males and females were used, and experiments were conducted with sex and age-matched mice. All experiments used littermate controls. The *LSL-Pole^P286R^* allele (a.k.a. B6.129S6-*Pole^tm1Dcas^*/J; Stock#037051) was previously generated and maintained in a pure C57BL/6J background ^21, 24^. Intestinal epithelial cell-specific expression of *Pole^P286R^*was achieved by crossing with *Vil-Cre* mice (a.k.a. B6.Cg-Tg(Vil1-cre)997Gum/J; Stock# 004586). Both strains are freely available from the Jackson Laboratory Repository. All studies were approved by the Institutional Animal Care and Use Committee (IACUC) and conducted in accordance with IACUC guidelines and the National Institutes of Health Guide for the Care and Use of Laboratory Animals.

### Induction of colitis

Colitis was induced in mice by administering 2.5 - 3% DSS (molecular mass 36-40 kDa; TdB Consultancy) for 5-7 days in drinking water ad libitum. DSS-water was then replaced by normal drinking water. For chronic colitis, mice received 2.5% DSS-water for 5 days, followed by a 2-week recovery on normal water. This DSS exposure and recovery cycles were repeated two additional times to yield three cycles of DSS-induced colitis.

### Histopathology and immunohistochemistry

Colons were washed with PBS, fixed in 4% paraformaldehyde, and embedded in paraffin. Whole colon tissue sections were stained with hematoxylin & eosin (H&E). Histopathological evaluation of colitis was made as a combined score for inflammatory cell infiltration (score, 0– 3), ulceration (score, 0–3), and area of crypt distortion (score, 0–3).

For immunohistochemistry, colon tissues sections were de-paraffinized and hydrated through decreasing concentrations of ethanol from 100% to 70%. For antigen retrieval, tissue sections were placed in a 95°C 10 mM sodium citrate solution (pH 6.0) for 20 minutes followed by a 30-minute cool down. Tissue sections were blocked using 5% goat serum for 1 hour, followed by incubation with primary antibody. After overnight incubation at 4°C, the tissue sections were washed three times and incubated with HRP-conjugated goat anti-rabbit IgG for 1 hour at room temperature. Following three washes, tissue sections were counterstained with hematoxylin. The images were taken using a KEYENCE BZ-X800 Microscope.

### Depletion of gram-negative bacteria

To deplete gram negative commensal bacterial load, neomycin (1g/L) was added into the drinking water. Mice were allowed to drink only neomycin solution adlib tum. To verify the gram-negative bacterial load depletion, fecal samples were collected, and fecal DNA was isolated by power fecal pro DNA kit (REF-51804). Then quantified abundance of gram-negative bacteria using the universal bacterial 16S rRNA gene by real-time qPCR.

### DNA and library preparation for Whole genome sequencing (WGS)

DNA was extracted from the tumor tissue with the QIAamp DNA Mini Kit (51306, Qiagen) and DNA concentration was measured by Qubit fluorometer (Invitrogen, Thermo Fisher Scientific). Sample integrity was confirmed by agarose gel electrophoresis. For preparation of libraries, 1.5 μg DNA was fragmented by Covaris ultrasonicator, then analyzed by gel electrophoresis.

The fragmented DNA was combined with End-Repair Mix and incubated at 20°C for 30 minutes. The end-repaired DNA was purified with QIAquick PCR Purification Kit (Qiagen) followed by addition of A-Tailing mix (Illumina) and incubated at 37°C for 30 minutes. This was combined with the purified Adenylated 3′-end DNA, adapter and ligation mix. The ligation reaction was incubated at 20°C for 15 minutes. Adapter-ligated DNA was run on a 2% agarose gel to recover target fragments. DNA was gel-purified with QIAquick Gel Extraction Kit (Qiagen). Several rounds of PCR amplification with PCR Primer Cocktail and Master Mix (both from Illumina) were performed to enrich Adapter-ligated DNA fragments. PCR products were run on a 2% agarose gel to recover the target fragments, followed by gel purification with QIAquick Gel Extraction Kit (Qiagen). The final library was analyzed by determination of average molecule length using an Agilent 2100 bioanalyzer (Agilent DNA 1000 Reagents) and quantification performed by real-time PCR by TaqMan assay. Qualified libraries were loaded onto the HiSeq X Ten sequencer (Illumina) for paired-end sequencing with read lengths of 100–150 bp.

### Mutation analysis

Sequencing reads were aligned to the mouse reference genome (GRCm38) using BWA 0.7.17 ^47^. Duplicate reads were marked using Picard, and base quality score recalibration was performed using GATK 4.2.0.0.0 ^48^. SNP and indel calling was conducted with Samtools ^49^.

Variants were considered present in a sample if the alternative allele frequency exceeded 0.1. Somatic mutations were defined as those absent from the Mouse Genomes Project database (including several FVB and 129 substrains) ^50^ and detected in only a single sample. All identified mutations were annotated with SnpEff ^51^, using GENCODE M16 annotations ^52^. Average sequencing depth was calculated for 1 Mbp windows using Samtools. Depth values were normalized to the genome-wide median and transformed using log2 scale. Base substitution rates were computed as the number of mutations divided by the number of positions covered by at least 20 reads. Trinucleotide mutational signatures were generated using a custom script.

### Murine colonic organoid culture

Colons collected from control and *Pole^P286R^* mutant mice were washed in PBS and cut into small pieces (2-3 mm) using sterile scissors. To isolate intestinal crypts, tissue pieces were resuspended in cell dissociation medium (STEMCELL) at room temperature for 15 minutes on a gently rocking shaker. The cell dissociation medium was removed, and the tissues placed into 2-8°C PBS. After pipetting several times, the supernatant was separated and filtered through 70μm cell strainer. These steps were repeated to collect up to 4 fractions. All fractions were centrifuged at 500 rpm and the tissue pellets resuspended in DMEM/F-12 media. The number of isolated crypts was counted under microscope and the volume adjusted to 500 crypts/ml. The tissue suspension was centrifuged again at 200*g* at 2-8°C for five minutes and the cell pellet was resuspended into 100 μl of complete InstestiCult^TM^ organoid growth medium (STEMCELL). Undiluted Matrigel matrix (100 μl) was added into the cell suspension and 20μl of this mixture containing 200 crypts was placed at the center of the well in a prewarmed 48-well cell culture plate. The plate was incubated at 37°C in a CO_2_ incubator for 10 minutes to allow the Matrigel to solidify. After the addition of 300 μl InstestiCult^TM^ organoid growth medium into each well, the plate was incubated at 37°C. The organoid growth medium was changed every four days.

### Statistical analysis

All the data were presented as means ± SD or ± SEM (as stated in the figure legends). Animal studies were repeated at least two times and in vitro studies were repeated 3 times. Statistical significance when comparing two groups was determined by two-tailed unpaired Student’s t-test. One-way ANOVA was used to analyze statistical significance between multiple groups. For statistical analysis of survival, Log-rank (Mantel Cox) test was used. p < 0.05 was considered statistically significant. GraphPad Prism 10.2.3 software was used for statistical analyses.

## ACKNOWLEDGEMNETS

We would like to thank the UT Southwestern Animal Resource Center (ARC) for maintenance and care of our mouse colony, and Deborah L. Gumucio for contributing the *Vil1-Cre* allele to the Jackson Laboratory. This work was supported by NIH 1R01CA265884 given to DHC and HZ. HZ acknowledges support from American Cancer Society (ACS) Research Scholar Grant (RSG-21-021-01-TBE), and NIH R01 DK125352 and R01DK128031 grants. We acknowledge the assistance of the UT Southwestern Tissue Resource Center, a shared resource at the Simmons Comprehensive Cancer Center, which is supported in part by the NCI under award number 5P30CA142543 for cutting, processing and H&E staining of tissue samples.

## AUTHOR CONTRIBUTIONS

DHC and HZ conceived and designed the study, analyzed data, and wrote the manuscript. MKM led the project, performed experiments, analyzed data, and wrote the initial manuscript. MA helped generate and maintain the mouse model, established the colitis protocol, and performed experiments. SR, MAB, and MA helped in experiments and data validation. HZ analyzed the exome sequencing data. EA shared reagents, contributed to the experimental plan and edited the manuscript.

## DATA AVAILABILITY

All raw data used in this paper will be submitted to journal as a part of publication and will be publicly available. *Pole^P286R^*mice are freely available at Jackson Laboratory. WGS data were deposited to NCBI and will be publicly available using the accession number PRJNA1367584.

## Abbreviations

CRC: colorectal cancer
DSS: dextran sulfate sodium
IBD: inflammatory bowel disease
LSL: Lox-Stop-Lox
POLE: DNA polymerase epsilon
POLD: DNA polymerase delta
PPAP: polymerase proofreading-associated polyposis
PVC: Pole^P286R/+^;Vil-Cre
WGS: whole genome sequencing
TMB: tumor mutation burden)

