## Supplemental text and Figures for "Inflammation is an essential trigger for POLE-driven ultramutated colorectal cancer"

**Supplemental material includes:**

1. Supplementary Methods
2. Supplementary Figures and Figure Legends (Figures S1-S6)
3. Supplementary Table 1

**SUPPLEMENTARY METHODS**

**Clinical assessment of colitis**

Body weight changes were monitored daily during each cycle of DSS treatment. Diarrhea scoring (0-3) was defined as follows: 0 = well-formed pellets, 1 = semi formed stools that did not adhere to the anus, 2 = semi formed stools that adhered to the anus, 3 = liquid stools that adhered to the anus. Bleeding scoring (0-3) was defined as follows: 0 = no blood by using hemoccult (Helena Pharmaceuticals), 1 = positive hemoccult, 2 = blood traces in stool visible, 3 = gross rectal bleeding.

**Real-time RT-PCR**

Colon tissues (normal colon or tumor) were frozen in dry ice immediately after being dissected from euthanized mice. Total RNA was extracted using Trizol (Invitrogen) according to the manufacturer’s instructions. Isolated RNA was reverse transcribed into cDNA using iScript (Bio-Rad). Real-time RT-PCR was performed using iTaq Universal SYBR Green Supermix (Bio-Rad). Expression data were normalized to GAPDH. Primers used for real-time PCR are listed in Supplementary Table 1.

**Western blot analyses**

Mouse normal colon tissues or colorectal tumors were collected and homogenized in RIPA lysis buffer containing complete protease and phosphatase inhibitor cocktail (Roche). Proteins were resolved by SDS-PAGE and transferred onto a polyvinylidene difluoride (PVDF) membrane. The membranes were immunoblotted with antibodies against Phospho-ERK (4370, Cell Signaling), ERK (4695, Cell Signaling), JNK, Phospho-JNK (4668, Cell Signaling), AKT, Phospho-AKT (4060, Cell Signaling), β-catenin (8480, Cell Signaling), Phospho-β-catenin (9561, Cell Signaling), GSK-3β (9315, Cell Signaling), Phospho-GSK-3βSer9 (9323, Cell Signaling), Phospho-NF-κB p65 (3033, Cell Signaling), cMyc (5605, Cell Signaling), Cyclin D1 (2978, Cell signaling), p53 (Sc98, Santa Cruz), P-Stat3 (9145, Cell Signaling), PCNA (13110, Cell Signaling), and β-actin (A2228, Sigma). Immunoreactive proteins were visualized by using ECL Super Signal West Femto substrate reagent (Thermo Scientific).

**Proliferation measurement using BrdU incorporation assay**

Proliferating cells were identified and counted with BrdU incorporation assay using BrdU *In-Situ* Detection Kit (550803, BD Pharmingen). BrdU (B5002, Sigma Aldrich) was administered (i.p.) into each mouse at a dose of 50mg/kg body weight 2 hours prior to euthanasia. Tissue samples were fixed in 4% paraformaldehyde and embedded in paraffin. The paraffin-embedded colon tissue sections were de-paraffinized by washing twice with xylene for 5 minutes each time at room temperature. The tissue sections were dehydrated by incubation in 100% ethanol 2 times for 5 minutes followed by once in 95% ethanol for 3 minutes at room temperature. The tissue sections were treated with 0.3% H_2_O_2_ to block endogenous peroxidase, followed by antigen retrieval with ‘BD Retrievagen A’ (#550803, BD Pharmingen) in a microwave oven at 89°C for 10 minutes. The tissue sections were incubated in biotinylated anti-BrdU antibody (#550803, BD Pharmingen) at 1:10 in diluent buffer for 1 hour at room temperature, and then in HRP-conjugated streptavidin for 1 hour at room temperature. The tissue sections were stained with DAB substrate solution followed by hematoxylin counterstaining. Slides were examined under microscope and the number of BrdU-positive cells per crypt counted.

**In vitro BrdU incorporation assay**

For measuring proliferation of organoid cells, organoid cultures were incubated with BrdU (10 μM) for 2 hours at 37°C with 5% CO_2._ The cells were then fixed in 70% ethanol for 5 minutes, treated with 1.5 N HCl for 30 minutes at room temperature followed by washing with PBS. After blocking with 5% normal goat serum in PBS and 0.3% Triton-X100 for 1 hour, the cells were incubated with anti-α-BrdU antibody (Bu20a, 5292, Cell Signaling) overnight at 4°C. After washing in PBS, the cells were treated with anti-mouse antibody conjugated with Alexa Fluor 488 for 2 hours at room temperature. Finally, the cells were washed with PBS and mounted with DAPI-containing mounting media.

**Immunostaining of organoids**

Organoids were grown in 8-well glass chamber slide with Intesticult organoid growth medium (mouse) supplemented with 50ug/ml Pen/Strep at 37°C and 5% CO_2_ incubator. Media was gently aspirated from the slide chamber and washed with PBS. Organoids were then fixed with 500ul of cold 4% PFA and left at room temperature (RT) for 30 minutes. After washing with PBS twice, Triton-X 100 (0.2% in PBS) was added and incubated at room temperature for 30 minutes. Organoid culture was washed with PBS three times and incubated with 5% BSA in PBS for 60 minutes at room temperature. After washing with PBS, organoids were incubated with primary antibodies at 4°C overnight. The culture was washed with PBS for five times and anti-rabbit HRP secondary antibody was added into each well. Following a 2 hour incubation at room temperature, organoids were washed with PBS and counterstained with DAPI. The stained organoids were observed under KEYENCE BZ-X800 Microscope.

**SUPPLEMENTARY FIGURES**

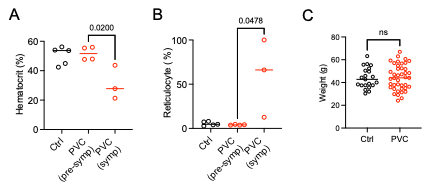

**Figure S1. *PVC mice small intestinal neoplasia*.** (A) Control and PVC mice was monitored for sickness. (A-B) Bloods collected from symptomatic and pre-symptomatic PVC mice were examined for percentage of Hematocrit (A) and Reticulocyte (B). (C) A separate cohort of control and PVC mice were monitored for survival. Body weights of control and PVC mice at euthanasia.

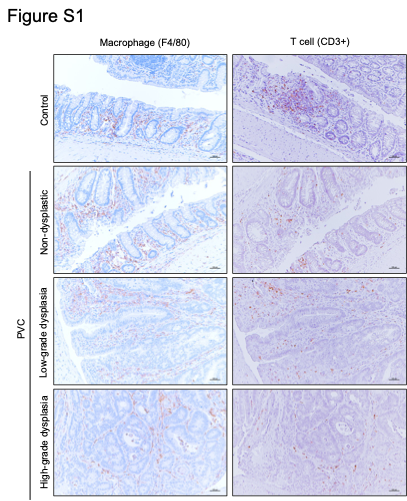

**Figure S2. *Pole^P286R^* tumors are infiltrated with macrophages and T cells**. Chronic colitis was induced in control and PVC mice with 3 cycles of 2.5% DSS regimen. Colons were collected 12 weeks post DSS treatment and processed for immunohistochemistry. Colon tissue sections were immunostained for macrophages and T cells with anti-F4/80 and anti-CD3 antibodies, respectively.

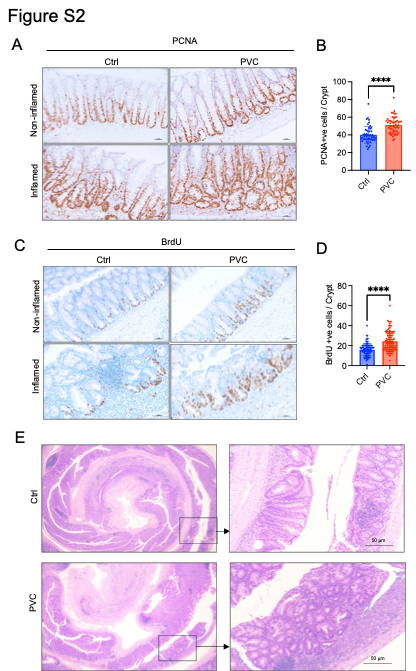

**Figure S3. Neoplastic lesions are developed in PVC mice during recovery phase of acute colitis.** Colitis was induced in control and PVC mice with the administration of 2.5% DSS for 5 days. Mice were sacrificed after 15 days of DSS administration and BrdU were administered (i.p.) 2 h prior euthanasia. (A) Colon tissue sections immunostained for PCNA. (B) The number of PCNA-positive cells per crypt. Data represent means ± SD (50 crypts per mouse, 2 mice per group), ****<0.0001, two-tailed unpaired t-test. (C) Tissue sections were stained with anti-BrdU antibody. (D) The number of BrdU-positive cells. Data represent means ± SD (50 crypts per mouse, 2 mice per group), ****<0.0001, two-tailed unpaired t-test. (E) H&E-stained colon tissue sections from control and PVC mice.

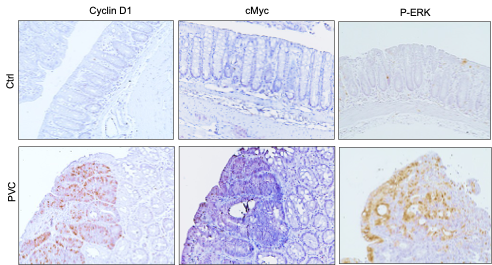

**Figure S4. Acute colitis activates oncogenic pathways in PVC mice.** Control and PVC mice were administered 2.5% DSS for 5 days. Colons collected at 4 weeks post DSS were examined for CyclinD1, cMyc and P-ERK with immunohistochemistry.

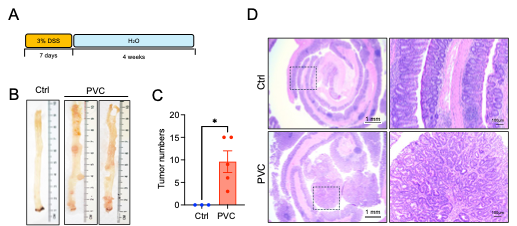

**Figure S5. A single DSS cycle is sufficient to induce adenocarcinoma in PVC mice.** (A**)** Control and PVC mice were administered with 3% DSS for 7 days. (B) Gross images of colons collected at 4 weeks post DSS. (C) Tumor burden in the colon. (D) H&E-stained colon sections. PVC mouse colons exhibited adenocarcinomas. Data represent means ± SEM, *P<0.05, **<0.01, two-tailed unpaired t-test.

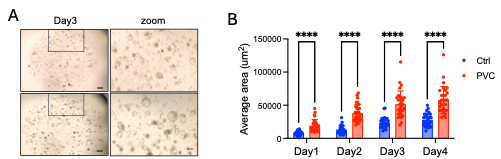

**Figure S6. *Pole^P286R^* mutation enhances growth of colonic organoids.** Colonic crypts from healthy (DSS untreated) control and PVC mice were cultured in 3D organoid culture system**.** (A) Organoid development was monitored microscopically. Images show organoids at day 3. (B) The size of organoids was measured at days 1, 2, 3 and 4. Data represent means ± SD, ****P<0.0001, two-tailed unpaired t-test.

**SUPPLEMENTARY TABLE 1.** Sequences of forward and reverse primers used for real-time RT-PCR analysis of indicated genes.

| Gene name | Forward | Reverse |
| --- | --- | --- |
| mIl1b | GCCTCGTGCTGTCGGACCCATA | TGCAGGGTGGGTGTGCCGTCTT |
| mIl6 | CAAGAAAGACAAAGCCAGAGTC | GAAATTGGGGTAGGAAGGAC |
| mTNFa | TCCCAGGTTCTCTTCAAGGGA | GGTGAGGAGCACGTAGTCGG |
| mIFNg | GAAAGACAATCAGGCCATCA | TTGCTGTTGCTGAAGAAGGT |
| mCxcl1 | TGAGCTGCGCTGTCAGTGCCT | AGAAGCCAGCGTTCACCAGA |
| mCxcl2 | CAAGAACATCCAGAGCTTGAGTGT | GCCCTTGAGAGTGGCTATGACTT |
| mKi67 | AGAAGTCCAGGTCTACAG | TCGTTGCTATTGCTAAGG |
| mMmp7 | ACCAACCTATTCCTGGTTGCTGCT | ATGGAAACGGGACAAGTCTGTGGA |
| mCtnnb1 | AAGGGCAAGGTTCGAATCAA | AGCCGAGATGGCCCAGAAT |
| mLgr5 | CGGGACCTTGAAGATTTCCT | GATTCGGATCAGCCAGCTAC |
| mMyc | CCTTTGGGCGTTGGAAACC | TCCTCGTCGCAGATGAAATAGG |
| mCcnd1 | TGCCATCCATGCGGAAA | AGCGGGAAGAACTCCTCTTC |
| mCdk4 | AATGTTGTACGGCTGATGGA | AGAAACTGACGCATTAGATCCT |
| mCcl2 | TTGCCGGCTGGAGCATCCACGT | AGTAGCAGCAGGTGAGTGGGGCG |
| mCdkn1a | TTGTCGCTGTCTTGCACTCT | ATCTGCGCTTGGAGTGATAG |
| mGapdh | TGGCAAAGTGGAGATTGTTGCC | AAGATGGTGATGGGCTTCCCG |
